# Organo-metal coprecipitation contributes to stable organic carbon fraction in mangrove soil

**DOI:** 10.1101/2025.02.15.638410

**Authors:** Kota Hamada, Nada Yimatsa, Toshiyuki Ohtsuka, Nobuhide Fujitake, Toshihiro Miyajima, Yusuke Yokoyama, Yosuke Miyairi, Morimaru Kida

## Abstract

Prediction of the impact of anthropogenic disturbances and global change on organic carbon (OC) pools in mangrove soils requires a detailed understanding of the mechanisms underlying OC stabilization. Using density fractionation to physically separate OC fractions with varying degrees of mineral association and protection, this study aimed to assess distributions of these fractions and the geochemical factors influencing the most dominant and refractory mineral-associated, high-density fraction (HF) in mangrove soil. We conducted forest-wide soil sampling in the Gaburumata mangrove forest on Ishigaki Island, Japan, along three transects (upstream, midstream, downstream) and at five depths (until 100 cm). The OC in HF (OC_HF_) was the oldest (median Δ^14^C value of -13.81‰) and contributed most significantly to total soil OC (43%-63%) and nitrogen (64%-85%). Among the extractable metals analyzed (aluminum [Al], iron [Fe], calcium [Ca], and magnesium [Mg]) with different crystallinity, only organically complexed Al and Fe showed strong positive correlations with OC_HF_. Together with high OC_HF_:Fe ratios that surpassed the maximum sorptive capacity of Fe oxides, these results indicate that co-precipitation of OC and Fe was the dominant mode of organo-mineral associations. The low clay content reduced the importance of Ca and Mg on OC_HF_, as these divalent cations typically facilitate OC stabilization through cation bridging between negatively charged clay surfaces and organic matter. Furthermore, the Δ^14^C–OC relationship suggested efficient incorporation of mangrove-derived modern C into HF, in addition to the pre-existing old C. Thus, mangrove expansion is likely to enhance stable soil OC pools in addition to increasing plant biomass and litter. Overall, this study proposes a biogeochemical mechanism for how stable mangrove OC is newly formed, as well as maintained, with ramifications for global mangrove expansion and plantation efforts.

## 1. Introduction

Vegetated coastal ecosystems such as mangroves, salt marshes, and seagrass meadows are vital carbon sinks, collectively termed "blue carbon” ecosystems (Mcleod et al., 2011). These ecosystems may accumulate soil organic carbon (OC) at rates several times higher than terrestrial ecosystems, offering a regional cost-effective and environmentally friendly approach to mitigate climate change and other adverse effects (Alongi, 2014; Macreadie et al., 2021; Mcleod et al., 2011). Among them, mangrove forests are particularly effective, sequestering OC at rates 4–5 times greater than terrestrial forests (Macreadie et al., 2021; Mcleod et al., 2011).

However, these ecosystems face significant threats from climate change and anthropogenic pressures, such as deforestation, land reclamation, and urbanization (Adame et al., 2021; Richards & Friess, 2016). Understanding the factors influencing OC stabilization in mangrove soils is critical for developing effective conservation and carbon management strategies.

Physical fractionation techniques, such as density and particle size separation, are widely used to investigate OC pools in soils. These methods distinguish particulate organic matter (POM), which is mainly plant-derived and exposed to microbial decomposition, from mineral-associated organic matter (MAOM), more microbially-processed, stable OC fraction with long turnover times (Heckman et al., 2021; Lavallee et al., 2020). Recent applications of these techniques and subsequent analysis of OC fractions in blue carbon ecosystems have provided deeper insights into OC composition, sources, protection mechanisms, and responses to environmental changes (Assavapanuvat et al., 2024; Broek et al., 2018; Fu et al., 2024; Hamada et al., 2024; Komada et al., 2022; Wang et al., 2022). For example, stable isotope analysis in a mangrove-salt marsh ecotone showed different contributions of contemporary (mangrove) and old (salt marsh) vegetation to POM and MAOM (Assavapanuvat et al., 2024). OC in mangrove forests originates from multiple sources, broadly classified as autochthonous (mangrove-derived) and allochthonous (terrestrial- or marine-derived) material (Bouillon et al., 2008). Fractionation of bulk OC into POM and MAOM helps disentangle different sources, while bulk analysis gives only their weighted mean. Density fractionation is particularly suitable for mangroves and salt marshes, as it can effectively separate and quantify root debris, which is a major autochthonous OC source in these ecosystems (Hamada et al., 2024; Komada et al., 2022). Radiocarbon dating has further revealed that MAOM represents the oldest and most stable OC pools in mangrove and salt marsh soils (Hamada et al., 2024; Komada et al., 2022). Yet, the mechanisms stabilizing OC in mangrove soils remain poorly understood, particularly the role of reactive minerals and metals under dynamic redox conditions (Kida & Fujitake, 2020).

Pedogenetic metals, particularly reactive aluminum (Al) and iron (Fe) (oxyhydr)oxides (e.g., amorphous, short-range ordered, and crystalline oxide minerals), enhance OC preservation through adsorption and co-precipitation, increasing the mean residence time of OC (Doetterl et al., 2015; Kaiser & Guggenberger, 2000; Oades, 1988). Co-precipitation of organic matter with Al/Fe ions as organo-metal complexes also contributes to its stabilization (Boudot et al., 1989; Takahashi & Dahlgren, 2016). However, in mangrove soils, the redox oscillations create a complex system where Fe can either stabilize OC in oxic conditions (Dicen et al., 2018; Lalonde et al., 2012; Li et al., 2024; Shields et al., 2016; Zhao et al., 2018) or destabilize it under anoxic conditions through reduction processes including Fe(II) dissolution, Fe respiration, and generation of reactive oxygen species under the presence of sulfide (C. Chen et al., 2020; Murphy et al., 2014; Thompson et al., 2006). Unlike Fe, Al, calcium (Ca), and magnesium (Mg) are less influenced by redox oscillations and thus may provide more stable OC associations under these conditions (Nitzsche et al., 2022; Rowley et al., 2017; Yu et al., 2021), but their roles in OC stabilization in mangrove soils are poorly understood. Furthermore, properties of the OC associated with these metals in mangrove soils, such as origin and age, remain unexplored.

To gain a deeper understanding of the preservation and accumulation of OC in mangrove soils, we conducted a forest-wide, comprehensive study in a mangrove forest on Ishigaki Island, Japan. Using density fractionation and subsequent analyses of stable and radiocarbon isotopes and reactive minerals and metals, we investigated the role of Fe, Al, Ca, and Mg in stabilizing OC in the mineral-associated fraction. Additionally, we conducted an analysis of endmembers that potentially serve as the origin of mangrove OC. We hypothesized that (i) organo-Fe co-precipitation would be important in determining OC content in the mineral-associated fraction in redox-dynamic mangrove soils, and (ii) OC incorporated by organo-Fe co-precipitation would show young age indicative of mangrove-derived origin.

## 2. Materials and Methods

### 2.1. Study area and sampling

The study area, located along the Gaburumata River on Ishigaki Island, Okinawa Prefecture, Japan (24°30’18.5"N, 124°14’52.2"E, Fig. 1), comprises a mangrove forest covering approximately 1.5 ha around the river estuary. Based on the field survey conducted for separate research, the vegetation is dominated by *Bruguiera gymnorrhiza*, with *Rhizophora stylosa* sparsely found only in the outer parts of the forest adjacent to creeks (Fig. 1c). We therefore considered that difference in vegetation was small and not a major factor influencing OC characteristics of different sampling locations. The climate of this region is subtropical monsoon, characterized by an average annual precipitation of 2116 mm and an average annual temperature of 24.6°C over the period from 1994 to 2023 (data from the Ishigaki Island Local Meteorological Observatory, Japan Meteorological Agency).

**Fig. 1.**
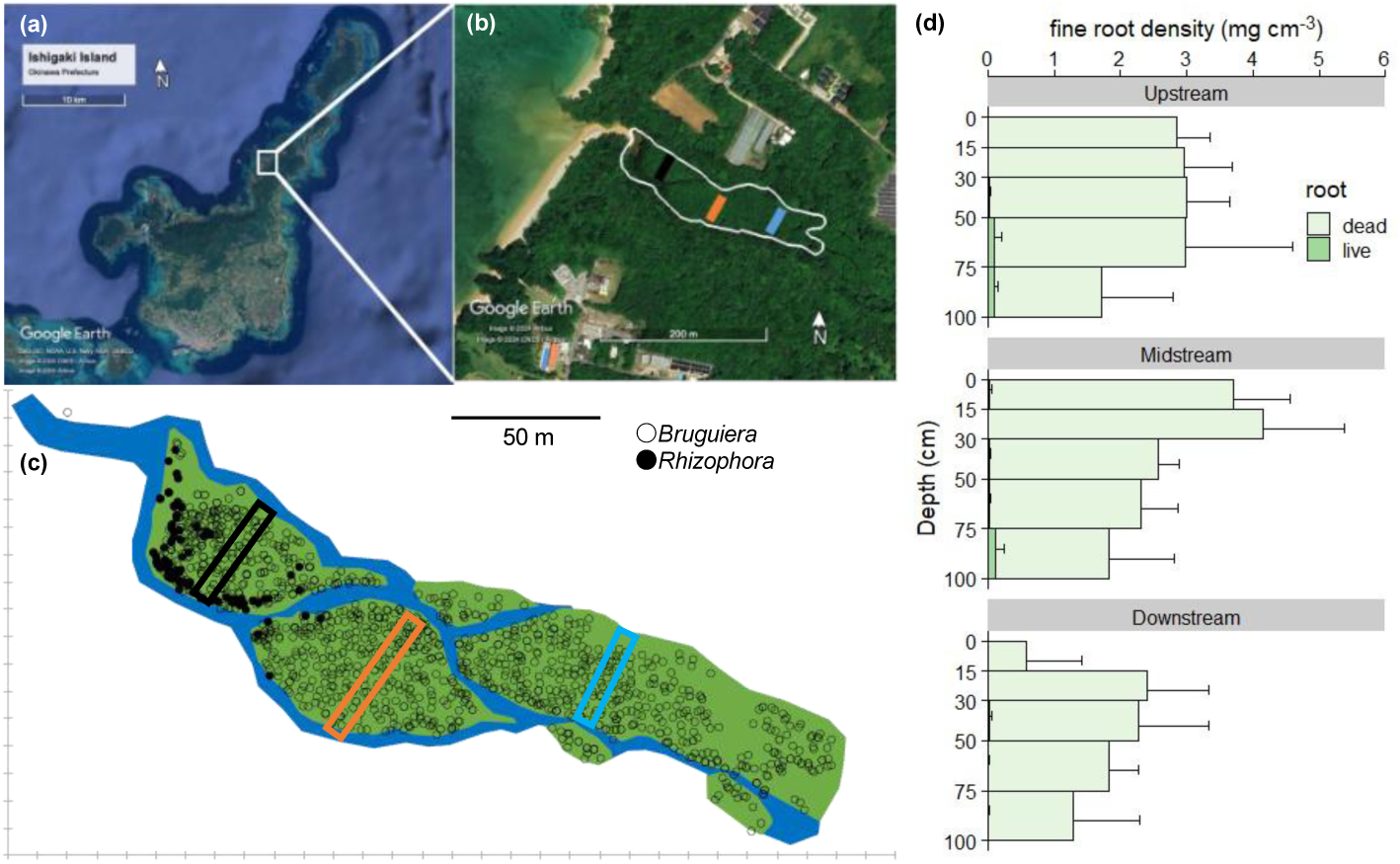
Map of the study site and fine root density. (a) Location of the Gaburumata River mangrove forest on Ishigaki Island, Japan. (b) The area enclosed by the white line represents the mangrove forest, and the bars within the area indicate the transects set for upstream (Blue), midstream (Orange), and downstream (Black) locations. (c) Detailed map of the Gaburumata River mangrove forest showing the sampling transects and all standing mangrove trees. The blue area indicates creeks. (d) Fine root density (mean + SD, *n* = 4) of each sampling location.

We established transects at three locations (upstream, midstream, downstream) to consider possible differences in soil properties by location (Fig. 1b, c). The elevations of three locations differed within only one meter and did not show clear patterns, nevertheless inundation time was presumably shorter at upstream given the delayed tidal movement. We collected soil samples in June 2022 at low tide at five depths: 0-10, 10-20, 20-40, 40-70, and 70-100 cm, respectively, using an open-face stainless-steel soil auger without compaction. We collected samples at ten points along each transect at approximately equal intervals and mixed corresponding depth sections to create one composite sample per depth per transect. We stored the composite samples in plastic bags and transported them to the laboratory under cool, dark conditions. The soil texture identified on-site was sandy. Composite samples were frozen and subjected to vacuum freeze-drying for seven days. Subsequently, we gently sieved freeze-dried soil through a 2 mm mesh sieve. Gravel was absent from all samples, but downstream samples contained white particles, likely shell fragments.

We also investigated fine root depth distribution at each transect by volumetric sampling using either the same open-face stainless-steel soil auger used for soil sampling (cross-section area of 27 cm^2^) or a Russian-type open-face peat sampler (9.8 cm^2^). Soil samples were collected up to a depth of 1 m near each transect in May 2024 (*n* = 4) and cut into similar depth intervals as soil samples (0-15, 15-30, 30-50, 50-75, and 75-100 cm). We washed off soil matrix at the field using creek water on a 0.5-mm mesh sieve and brought roots back to the laboratory where they were stored in a refrigerator. Then, we washed them again with tap water on a 0.5-mm mesh sieve. All roots remaining on the sieve were manually sorted into four categories in water based on diameter (fine root, < 2 mm; coarse root, ≥ 2 mm) and vitality (living or dead), which was evaluated by their color and firmness (Poungparn et al., 2015).

Living roots were identified by their whitish or brownish color, resilience, firmness, and ability to float in fresh water. All roots were oven-dried at 70 °C for over 48 hours and weighed. Root density distributions were estimated based on the section volume and root weight. Because of the low and patchy abundance of coarse roots, we present only fine root distributions (Fig. 1d).

As endmembers potentially contributing to OC in the mangrove soils, we collected the following endmember samples: (1) terrestrial soil samples (A1 and A2 layers) from the catchment of the Gaburumata River, (2) riverbank soil (0–10 cm) of the headwater of the Gaburumata River, (3) riverine suspended solids (SS) at the nearby Fukido River headwater, collected using a suction filtration device with a 0.45 µm PTFE membrane filter (Omnipore, Merck, Germany), (4) green leaves and fine roots of *Bruguiera gymnorrhiza* and *Rhizophora stylosa* from multiple standing trees, (5) senescent yellow leaves of *Bruguiera gymnorrhiza* found on the ground within the forest, (6) seagrass in the estuary of the Fukido River, (7) sandy soil beneath the seagrass meadow (*Halodule* sp., composites of triplicate cores by two depths, 0–5 and 5–10 cm), (8) macroalgae in the estuary of the Fukido River, and (9) marine SS in the open ocean north of Ishigaki Island, collected on a combusted, pre-weighed 0.7 μm glass fiber filter (Whatman GF/F, Cytiva, USA). Details of these samples are provided in Table 1. We assumed that the characteristics of seagrass, macroalgae, and SS samples from around the Fukido River (24°29’7’’N, 124°13’50’’ E) are similar to those contributing to the Gaburumata River, given the proximity of the two rivers (only ∼ 2 km away) flowing into the same coastal area and neighboring catchments with highly similar land use (mostly broadleaf forest) (Kida, Tanabe, et al., 2019). All samples were transported back to the laboratory under cool, dark conditions. Soil samples were frozen, freeze-dried, and sieved through a 2 mm mesh sieve and ground using a vibrating mill. Biomass samples were dried at 60°C and ground either with a vibrating mill or using a pestle and mortar. The riverine SS sample on a PTFE filter was recovered with pure water, freeze-dried, and ground, while marine SS samples were directly analyzed for elemental and isotopic composition after oven drying at 60°C.

**Table 1.** Organic carbon content (OC, mg C g^-1^), total nitrogen content (TN, mg N g^-1^), C/N ratio, and δ^13^C stable isotope ratios (‰) for various endmembers (mean ± SD).

| | Sampling year | Depth (cm) | OC (mg C g<sup>-1</sup>) | TN (mg N g<sup>-1</sup>) | C/N | $\delta^{13}\text{C}$ (‰) | *n* |
| --- | --- | --- | --- | --- | --- | --- | --- |
| Terrestrial soil (A1) | Jan 2014 | 0–7 | 52 | 3.1 | 17.3 | -28.14 | 1 |
| Terrestrial soil (A2) | Jan 2014 | 7–16 | 20 | 1.2 | 16.7 | -28.41 | 1 |
| Riverbank soil | June 2022 | 0–10 | 13 | 1.2 | 10.8 | -28.14 | 1 |
| Riverine suspended solid | Dec 2022 | - | 146 | 8.6 | 17.1 | -30.42 | 1 |
| Green leaf (*B. gymnorrhiza*) | June 2022 | - | 439 ± 11 | 12.4 ± 1.7 | 36.0 ± 4.9 | -33.71<sup>†</sup> | 4 |
| Yellow leaf (*B. gymnorrhiza*) | June 2022 | - | 433 ± 3 | 3.0 ± 0.2 | 146 ± 6.8 | -30.98<sup>†</sup> | 6 |
| Green leaf (*R. stylosa*) | June 2022 | - | 452 ± 13 | 8.1 ± 6.6 | 97.1 ± 72.7 | -30.60<sup>†</sup> | 4 |
| Fine root (*B. gymnorrhiza*) | June 2022 | - | 405 ± 24 | 7.8 ± 0.6 | 52.3 ± 4.3 | -28.76<sup>†</sup> | 4 |
| Fine root (*R. stylosa*) | June 2022 | - | 367 ± 25 | 6.8 ± 0.6 | 54.1 ± 2.3 | -28.47<sup>†</sup> | 4 |
| Seagrass meadow soil | June 2022 | 0–5 | 3.6 | 0.4 | 9.0 | -22.57 | 1<sup>‡</sup> |
| Seagrass meadow soil | June 2022 | 5–10 | 4.4 | 0.5 | 8.8 | no data | 1<sup>‡</sup> |
| Seagrass | May or Dec 2014 | - | 292 ± 46 | 23.9 ± 4.1 | 14.7 ± 3.7 | -9.32 ± 1.56 | 22 |
| Macroalgae | May or Dec 2014 | - | 193 ± 63 | 14.1 ± 6.1 | 19.1 ± 9.7 | -13.50 ± 2.68 | 20 |
| Marine suspended solid | July 2012 | - | 62 ± 24 | 12.5 ± 4.6 | 5.8 ± 0.3 | -22.35 ± 0.60 | 20 |

<sup>†</sup> Composite samples were analyzed.
<sup>‡</sup> Composite of triplicate cores.
Seagrass: *Cymodocea rotundata*, *C. serrulate*, *Enhalus acoroides*, *Halophila ovalis*, *Thalassia hemprichii*, *Halodule* sp.
Macroalgae: *Padina minor*, *Sargassum* sp., *Acetabularia ryukyuensis*, *Caulerpa* sp., *Ulva* sp., *Codium intricatum*.

### 2.2. Density fractionation

#### 2.2.1. Optimization of experimental conditions

We performed density fractionation using an aqueous solution of sodium polytungstate (SPT-0, TC-Tungsten Compounds, Germany; SPT) with a density of 1.6 g cm^-3^, following an established method (Hamada et al., 2024). Density fractionation separates total OC into free low-density fraction (f-LF), mineral-associated LF (m-LF), and high-density fraction (HF), where LFs are conceptually and operationally similar to POM and HF to MAOM, respectively. This method, widely used in terrestrial soils to separate soil organic matter fractions of varying microbial degradability, has seen limited application in mangrove soils (Assavapanuvat et al., 2024; Hamada et al., 2024). Given the unique properties of wet soils such as mangroves, optimizing density fractionation conditions is crucial. We investigated the effects of soil sample conditions (field-moist versus freeze-dried) and appropriate ultrasonic output (20 W, 40 W, 60 W, 80 W) during m-LF recovery, conditions we could not check in a previous study (Hamada et al., 2024). Detailed designs and results are presented in Supplementary Information (S1, Figs S1, S2). Briefly speaking, we found that freeze-dried samples produced results highly comparable to field-moist samples in relative distributions of density fractions while offering benefits of improved sample homogeneity and preservability (Fig. S1). We also found that m-LF mass recovery was largely unaffected by ultrasonic output, while processing at 20 W was the most efficient due to much reduced time needed for rinsing m-LF (Fig. S2). We thus conducted subsequent density fractionation under these optimized conditions (120 J mL^-1^, 20 W, freeze-dried soils).

#### 2.2.2. Sample analysis

To conduct density fractionation on the main samples, we weighed 5 g of freeze-dried soil in a 50 mL conical tube. Subsequently, we added 20 mL of SPT solution and gently inverted the tube 20 times. We then centrifuged the solution at 2800 G for 5 minutes and collected the suspended materials (f-LF) using a poly dropper and dispensing spoon onto a 0.45 µm PTFE membrane filter placed on a suction filtration device (Omnipore, Merck, Germany). This process was repeated twice, and after each collection, any lost SPT solution was replenished. To prevent potential inorganic nitrogen contamination from the SPT solution, we washed the collected f-LF three times with 5 mL of 1M KCl, followed by rinsing with deionized water until the electrical conductivity (EC) reached below 50 μS cm^-1^ and freeze-drying. Subsequently, we adjusted the volume of the remaining SPT solution in the conical tube to the 30 mL mark and treated the suspension with sonication under the optimized conditions described in Supporting Information (120 J mL^-1^, 20 W, 3 minutes). We centrifuged the sample at 12400 G for 20 minutes and carefully recovered the floating material (m-LF) by decantation. We rinsed the collected m-LF with a KCl solution and water in the same manner as in f-LF. Finally, we transferred the residue (HF) in the conical tube to a 250 mL centrifuge tube with deionized water and centrifuged at 14000 G for 25 minutes. We then carefully discarded the supernatant using a Komagome pipette. We treated the HF samples similarly as for LFs: rinsing with 100 mL of 1M KCl, shaking reciprocally for 10 minutes, centrifuging at 14000 G for 25 minutes, and washing several times with 100 mL of deionized water until the EC dropped below 50 μS cm^-1^, followed by freeze-drying. Finally, we measured the mass of each fraction and calculated the mass recovery. We ground the HF and LF samples using a vibrating mill and with a pestle and mortar, respectively.

### 2.3. Elemental analysis

We determined the elemental composition of bulk soil, each density fraction, and endmember samples using an elemental analyzer (Vario EL cube, Elementar, Germany). Bubbling was observed upon addition of 1M HCl to random soil samples at each location, highlighting the need for inorganic carbon (IC) removal prior to elemental analysis. We employed the vaporous method (Komada et al., 2008), where soil samples encapsulated in Ag capsules were exposed to vaporized concentrated HCl to remove IC (details in S2.1). After IC removal, we performed elemental analysis in duplicate, and the average values are presented. The analytical precision for C and N, defined as the absolute difference of element content between duplicate analysis, was on average 0.06% and 0.004%, and never exceeded 0.57% and 0.04%, respectively. From the obtained C and N content, we calculated mass recovery of C and N by density fractionation, contribution of each density fraction to bulk C and N content, and C/N mass ratios.

### 2.4. Stable carbon isotope analysis

We determined stable carbon isotope ratios of each density fraction and endmember samples using a continuous flow elemental analyzer/isotope ratio mass spectrometer (EA/IRMS; FLASH EA 1112 series + Thermo Finngan DELTA plus, Thermo Scientific, USA). The analysis was conducted after IC removal, either by HCl fumigation or by a drop of 1M HCl in Ag capsules (seagrass and macroalgae samples). We report carbon stable isotope ratios in delta (δ) notation as the per mil (‰) difference in the ^13^C/^12^C ratio relative to the Vienna Pee Dee Belemnite (VPDB) standard. Calibration standards L-Histidine (δ^13^C: -11.4‰ ± 0.2‰) and Glycine (δ^13^C: -32.3‰ ± 0.2‰) from Shoko Science, Japan, were utilized. To correct deviations from true values, we performed calibration with L-Histidine and Glycine standards at approximately 0.04 mg and 0.03 mg, respectively (Hamada et al., 2024). Isotopic ratios were calibrated and normalized to the international scale using a 5-point calibration method with these standards. Quality control measures included running two standards per every 20 samples, with blanks and conditioning/calibration standards at the beginning and end of each run. Analytical precision, defined as the absolute difference of δ^13^C values between duplicate analysis, was on average 0.31‰.

### 2.5. Radiocarbon analysis

We conducted radiocarbon analysis of density fractions and endmembers at Yokoyama Lab, AORI, Japan (Yokoyama et al., 2019). Approximately 1.5-3 mg of OC was provided for analysis, following IC removal as previously described. Samples were sealed in Ag capsules and combusted to CO_2_ gas using an elemental analyzer (Vario Micro Cube, Elementar), followed by conversion to graphite using a custom-built graphitization vacuum line (Yokoyama et al., 2022). Graphitized samples underwent radiocarbon measurement using single-stage accelerator mass spectrometry(SSAMS; NEC, USA). Results are reported in Δ^14^C, representing the fractional deviation (in parts per thousand, ‰) of the sample’s ^14^C/^12^C ratio relative to that of the oxalic acid international standard (National Institute of Standards and Technology) (Stuiver & Polach, 1977). Positive Δ^14^C values indicate the presence of bomb-^14^C derived from atmospheric nuclear weapons testing from 1959 to 1963, whereas negative values indicate the apparent residence time of the soil C. We considered all possible mass fractionations by measuring δ^13^C values using the AMS instead of using δ^13^C values measured in section 2.4. Analytical precision for Δ^14^C analysis was on average 2.4‰ and never exceeded 3.5‰.

Where necessary, the ^14^C data were converted into apparent calendar age using radiocarbon chronology (calAD) using OxCal 4.4 (Ramsey, 2009) with the IntCal20 NH3 curve (Reimer et al., 2020) for pre-bomb samples and Bomb21 NH3 curve (Q. Hua et al., 2022) for post-bomb samples. The median calAD years of the 2σ ranges (95.4% probability ranges) are presented to indicate the years of the photosynthetic assimilation of the measured C. All LF samples and some HF samples exhibited “modern” ^14^C ages with positive Δ^14^C values, where the calibration curve produced two possible calendar age ranges before and after the bomb peak. With an assumption that LF were mangrove-derived litter with a rapid turnover time and HF were older than LF, we assigned younger ages (i.e., calibration by the right side of the bomb peak) to LF and older ages (i.e., calibration by the left side of the bomb peak) to HF. When ^14^C ages of LF samples were so young that they were partially out of the calibration range (up to 2019 calAD), the most conservative (young) calendar year was assigned to the maximum probability range, which was the year of sampling (2022). When multiple calAD ranges were possible in pre-bomb HF samples, the probability-weighted mean of multiple calAD years was calculated for the 2σ range and median.

### 2.6. Extractable metals analysis

To extract reactive Al and Fe from HF samples, we employed standard methods including 0.75 M sodium dithionite-citrate extraction and 0.1 M sodium pyrophosphate extraction (Holmgren, 1967; McKeague, 1967). A sodium dithionite-citrate solution extracts metals through reductive dissolution of mineral phases via the reducing ability of dithionite, followed by complexation with citrate ligands, and thus conceptually dissolves all redox-sensitive elements in a soil (Holmgren, 1967; Mehra & Jackson, 1958; Parfitt & Childs, 1988). Citrate facilitates the removal of alumina coatings from the soil matrix (Mehra & Jackson, 1958). The metals extracted by sodium dithionite-citrate are referred to as reactive minerals and metals, which include organo-Fe complexes, iron sulfide, and Fe oxides including ferrihydrite, hematite, and goethite (but not pyrite), and Al and manganese oxides associated with Fe oxides. A sodium pyrophosphate solution at pH 10 is a weaker extractant and primarily dissolves metals from organo-metal complexes (i.e. polyvalent cations such as Al and Fe ions complexed with organic ligands containing carboxylic and phenolic groups) through complexation with pyrophosphate ligands (McKeague, 1967; Parfitt & Childs, 1988).

Briefly, each extraction of Al and Fe (sodium dithionite-citrate extraction and sodium pyrophosphate extraction) was conducted with a solid-to-liquid ratio of 1:100 and reciprocal shaking at room temperature for 16 hours. After extraction and high-speed centrifugation (details in S2.2), we measured the concentrations of Al and Fe in the extracts using inductively coupled plasma atomic emission spectroscopy (ICP-AES, SPS3100 series, High Tech Science Corporation, Japan) by the internal standard method with calibration curves matching the sample matrix. The Al and Fe extracted via dithionite-citrate reductive dissolution and sodium pyrophosphate were denoted as Al_d_ and Fe_d_, and Al_p_ and Fe_p_, representing all reactive minerals and organo-metal complexes through co-precipitation, respectively. Their differences (Al_d-p_, Fe_d-p_) are approximations of the sum of crystalline and nano-crystalline Al and Fe minerals, respectively (Parfitt & Childs, 1988).

We extracted exchangeable Ca (Ca_ex_) and Mg (Mg_ex_) using a modified batch method involving 0.05 M ammonium acetate and 0.0114 M strontium chloride (SrCl_2_) with a solid-to-liquid ratio of 1:200 and 1-h reciprocal shaking at room temperature, a technique widely applied in Japan (Kamewada & Shibata, 1997; Matsue & Wada, 1985) (details in S2.2). After extraction and high-speed centrifugation, we spiked the supernatant with 250 ppm yttrium and analyzed it by ICP-AES employing the internal standard method with calibration curves matching the sample matrix. Seawater-derived Ca and Mg did not interfere with our analysis, as the washing steps in density fractionation effectively removed sea salts. However, high sodium concentration of SPT (∼1.5 M) presumably displaced certain percentages of exchangeable Ca and Mg from the soil matrix. Consequently, exchangeable Ca and Mg measured in this study, extracted primarily by Sr, represent fractions strongly bound to the soil matrix.

### 2.7. Statistical analysis

We analyzed correlations between the contents of extractable metals and OC_HF_ by the ordinary least square method in *R* (https://www.r-project.org/). We assessed differences in Δ^14^C values between different fractions by Kruskal-Wallis rank sum test for three-group comparison and Wilcoxon rank sum test for two-group comparison. P-values lower than 0.05 were considered significant.

## 3. Results

### 3.1 Fine root distribution and density fraction abundance

Fine root density was similar irrespective of locations (Fig. 1d). Dead fine roots were dominant and accounted for 99% of the total fine root density. Fine roots were abundant until a depth of 1 m and their depth distributions were not clear without consistent patterns among locations (Fig. 1d).

The recovery of mass after density fractionation averaged 95.4%±7.6% (Table 2). Similarly, C and N recovery averaged 94.4%±5.5% across samples (Table 2). The C and N contents in LF were substantially higher compared to in HF, with C and N levels in LF being 20 and 15 times greater than those in HF, respectively (Fig. S3bc). Among LFs, m-LF consistently displayed higher C and N contents than f-LF (Fig. S3bc). Bulk soil-based abundance of LF as C (mgC g soil^-1^) showed an increasing trend with depth (Fig. 2a–c). HF contributed most to total soil C (43%-62%) and N (63%-81%), followed by f-LF (17%-38% of C and 9%-22% of N) and m-LF (16%-23% of C and 10%-18% of N), respectively (Fig. 2cd).

**Fig. 2.**
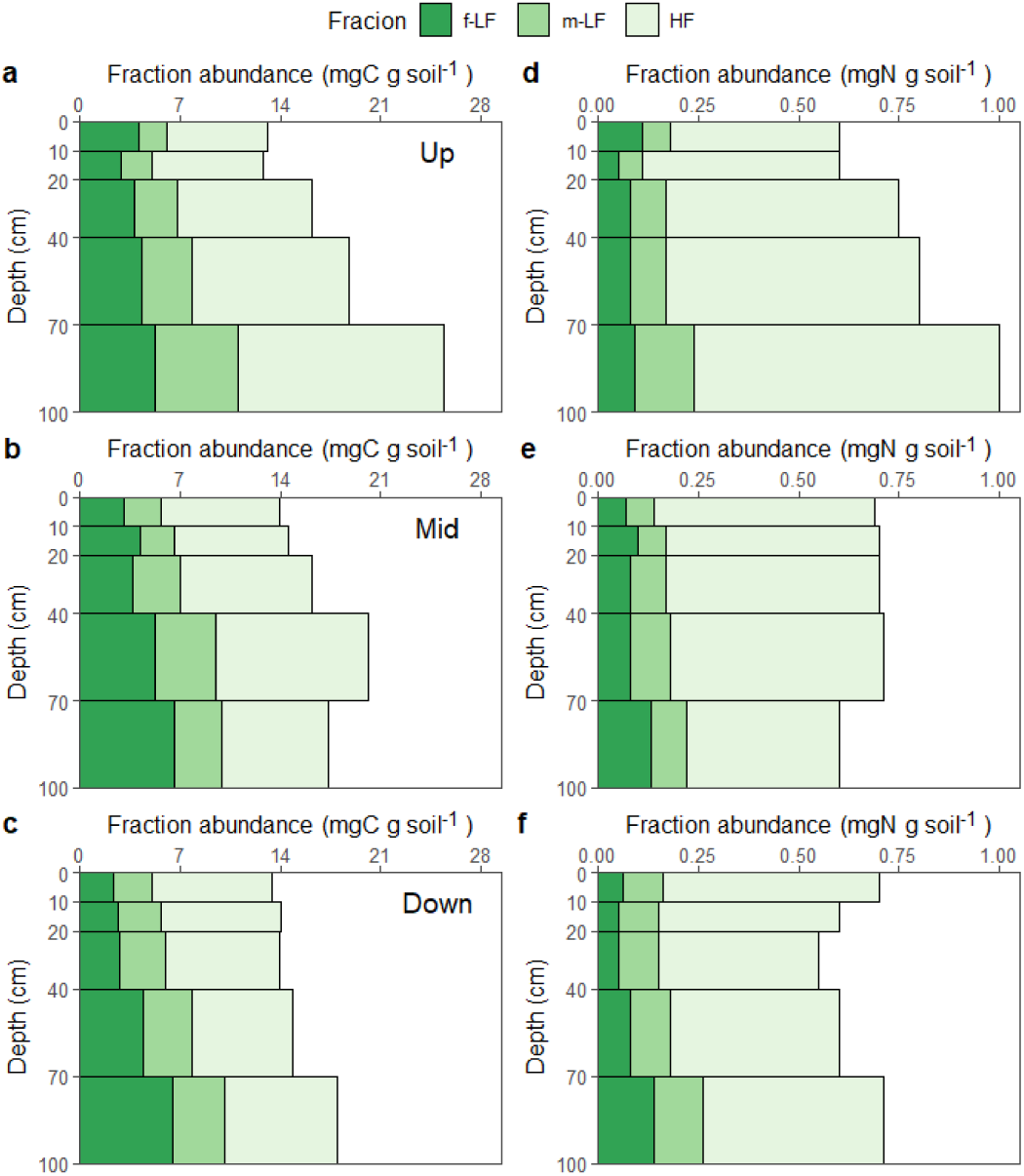
Contribution of each density fraction to total soil organic C (a–c) and N (d–f) content with depth. The upper (a, d), mid (b, e), and lower (c, f) panels are results for sampling locations at upper-, mid-, and downstream. Each data represents a composite sample of 10 different sampling (also in other figures). f-LF: free low-density fraction, m-LF: mineral-associated low-density fraction, and HF: high-density fraction.

**Table 2.**
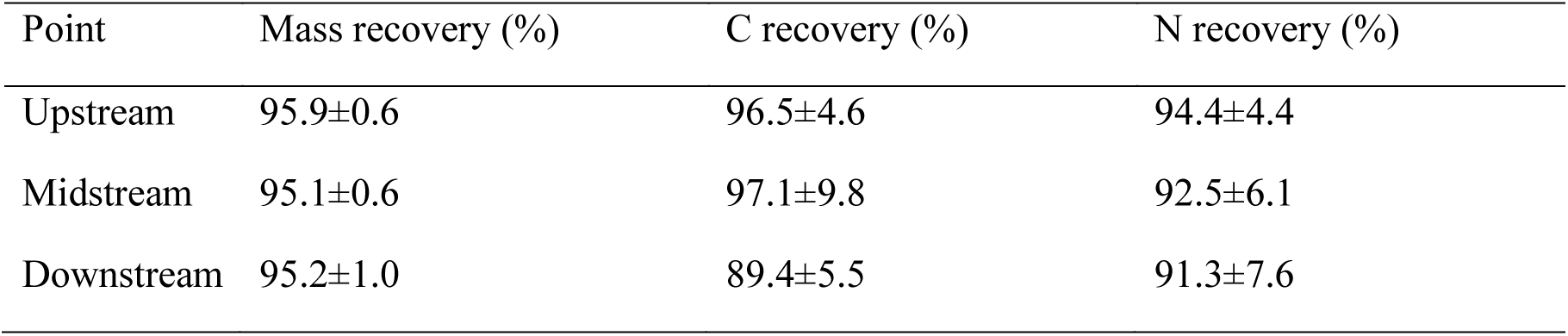
Mass and CN recovery of density fractionation.

| Point | Mass recovery (%) | C recovery (%) | N recovery (%) |
| --- | --- | --- | --- |
| Upstream | 95.9±0.6 | 96.5±4.6 | 94.4±4.4 |
| Midstream | 95.1±0.6 | 97.1±9.8 | 92.5±6.1 |
| Downstream | 95.2±1.0 | 89.4±5.5 | 91.3±7.6 |

### 3.2 C/N ratios and δ^13^C values

The carbon-to-nitrogen (C/N) mass ratios ranged from 36.3 to 65.1 for f-LF, 24.7 to 41.9 for m-LF, and 14.8 to 22.1 for HF (Fig. 3a). Generally, the C/N ratio followed the order f-LF > m-LF > HF, with all fractions showing an increase in C/N ratio with depth (Fig. 3a). The δ^13^C values of the density fractions were as follows: f-LF ranged from -29.7‰ to -28.6‰, m-LF from - 30.1‰ to -28.7‰, and HF from -29.1‰ to -27.9‰ (Fig. 3b). These low δ^13^C values across all samples indicated that terrestrial C3 plants, including mangroves, were the primary source of OC in this mangrove forest. HF displayed slightly elevated δ^13^C values than f-LF and m-LF, with downstream HF samples showing consistently higher δ^13^C values compared to those from mid- and upstream locations (Fig. 3b). A consistent trend of increasing δ^13^C values with depth was observed across all fractions (Fig. 3b).

**Fig. 3.**
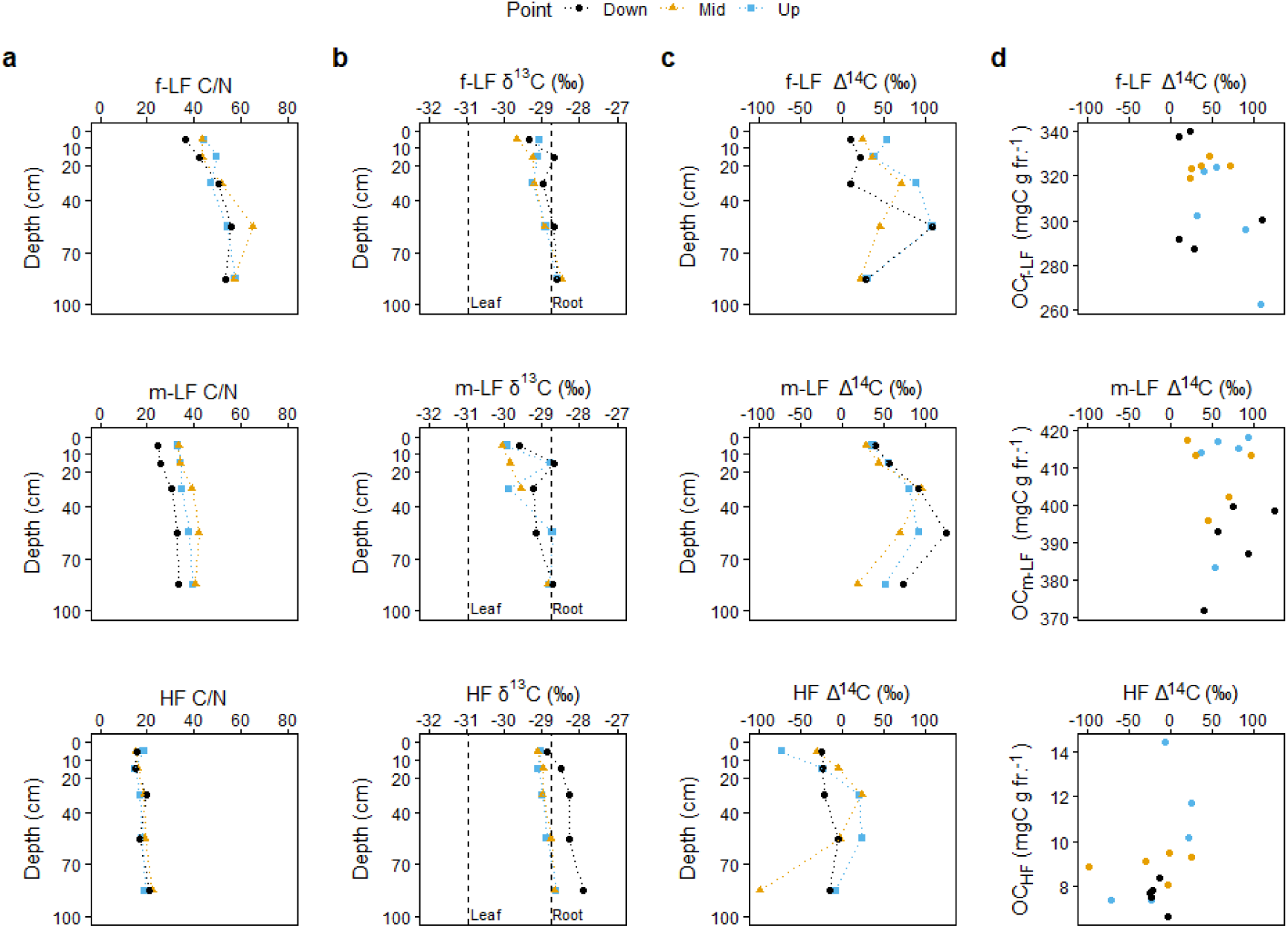
C/N ratio (a), δ^13^C stable isotope ratio (b), and Δ^14^C value with depth (c) and versus OC contents (d) of each fraction. The vertical dashed lines in (b) represent the average δ^13^C values for senescent, yellow leaf and fine root of mangrove (*Bruguiera gymnorrhiza*).

Endmember values exhibited high variability (Table 1). Large differences in C/N ratios were observed between the fine roots and green leaves of both *Bruguiera gymnorrhiza* and *Rhizophora stylosa* (Table 1). Senescent leaves of *Bruguiera gymnorrhiza* exhibited exceptionally high C/N ratios, indicating preferential N loss during initial decomposition or N translocation prior to leaf fall. Compared to mangrove samples, soil and suspended solid (SS) samples had much lower C/N ratios irrespective of terrestrial or marine origin (Table 1).

Seagrass and macroalgae also had low C/N ratios. Mangrove roots consistently showed higher δ^13^C values than corresponding leaves (Table 1). The highest δ^13^C values were recorded in marine primary producers (seagrass and macroalgae), followed by marine SS, seagrass meadow soil, terrestrial soil, mangrove roots, riverine SS, and mangrove leaves (Table 1). The substantial OC content and lower δ^13^C values of riverine SS compared to terrestrial soils indicate enrichment of C3 plant residues.

### 3.3 Radiocarbon

We observed clear differences in Δ^14^C values between HF and LFs across all measured cores (Fig. 3c). HF was consistently older (Kruskal-Wallis, p < 0.001) with Δ^14^C values between -

98.22‰ and 24.14‰, while all f-LF (9.87–108.71‰) and m-LF (18.9–124.69‰) samples exhibited modern values (young, decades-old C) with positive Δ^14^C values indicating an incorporation of a significant proportion of the bomb-^14^C (Fig. 3c). Although not significant (Wilcoxon, p = 0.08), a slight increase in Δ^14^C values in m-LF (median = 56.41‰) compared to f-LF (median = 36.28‰) was observed (Fig. 3c). These Δ^14^C values of LFs suggest slightly older ages of m-LF than f-LF assuming that LFs originated from photosynthetic production postdating the 1960s bomb peak.

The depth profile of Δ^14^C values of density fractions exhibited very similar patters irrespective of locations (Fig. 3c). Consequently, Δ^14^C values showed relatively high correlations between fractions (f-LF vs m-LF: R = 0.66, p < 0.01; f-LF vs HF: R = 0.53, p < 0.05; m-LF vs HF: R = 0.69, p < 0.01), suggesting interconnected C dynamics among these fractions. All fractions showed a Δ^14^C peak at mid depths (20–40 cm or 40–70 cm), with HF showing a transition from negative to positive Δ^14^C values, indicating considerable fresh carbon input to HF at these depths (Fig. 3c). This increase in Δ^14^C values were accompanied with higher OC content in HF (Fig. 3d), suggesting a build-up of modern OC on the pre-existing old OC. Contrarily, the increase in positive Δ^14^C values of f-LF and m-LF likely reflected longer turnover times of these fractions at these depths.

The timescales of OC dynamics among density fractions at different depths became clearer when calibrated to calendar years (calAD) (Fig. 4). Please note that they are apparent ages and are not indicative of real dates since samples are mixtures of different compounds. The calAD indicates the year of carbon fixation due to photosynthesis of each sample. f-LF and m-LF were oldest (∼21 years) at mid depths (20–40 cm or 40–70 cm) and younger (∼12 years) at shallower or deeper depths (Fig. 4ab). f-LF was on average 1–9 years older than m-LF at corresponding depth (Fig. 4ab), corroborating Δ^14^C results (Fig. 3c). The root density data, however, did not align with this depth distribution of LFs ages, as roots (i.e., modern C source) were not more abundant at the surface or bottom sections than the mid sections (Fig. 1d). The age distribution of HF was more irregular compared to that of LF (Fig. 4c). Broadly speaking, HF tended to be younger at mid depths and older at the surface or bottom, an opposite trend to LF (Fig. 4). However, a location difference was observed, with the oldest age found at the surface at the upstream location (weighted calAD of 1428) while at the deepest section at the midstream location (weighted calAD of 1266). At the downstream location, HF samples had less variable calAD years (1825–1955) (Fig. 4c).

**Fig. 4.**
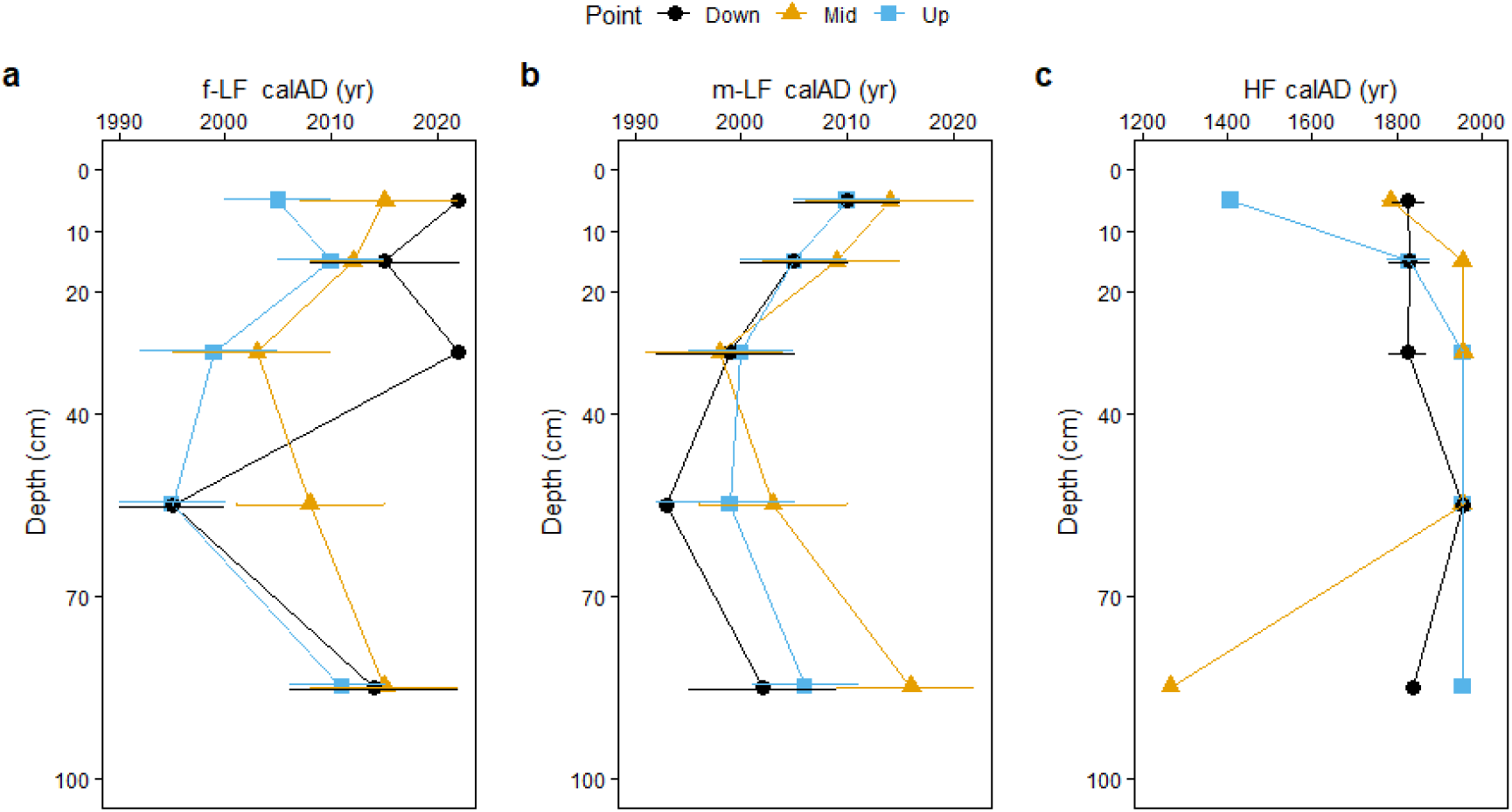
Apparent calendar years (year of carbon fixation due to photosynthesis) of density fractions along depth. The error bars indicate the minimum and maximum estimate of calAD (2σ range), with points representing the median. Error bars may be smaller than points in HF (c). Note the difference in scale of the x-axis between (a, b) and (c). The year of sampling was in 2022.

Endmember samples exhibited variable calAD years: terrestrial soil A1, 2004; terrestrial soil A2, 2003; riverbank soil, 1953, and riverine SS, 1827 calAD. The calendar year of riverine SS was close to that of some of HF than to LFs (Fig. 4).

### 3.4 Extractable metals (Al, Fe, Ca, Mg)

Reactive metals (Al and Fe) of varying crystallinity showed diverging patterns (Fig. S4). The dithionite-citrate extractable Al and Fe (Al_d_ and Fe_d_, representing all reactive minerals) contents decreased sharply at the surface (Fig. S4a, d), while pyrophosphate extractable fractions (Al_p_ and Fe_p_, representing organically complexed metals) did not display such abrupt change (Fig. S4b, e). Across all locations and fractions, Fe_d_ consistently exceeded Al_d_ by a factor of approximately 3 to 7 (Fig. S4a, d), whereas Al_p_ and Fe_p_ contents were roughly similar (Fig. S4b, e). Contrarily, when comparing distributional patterns of Al and Fe, these metals showed strong correlations (r > 0.84, p < 0.001) irrespective of crystallinity (not shown), indicating a shared mechanism controlling their abundance. The high levels of Fe_d_ resulted in a similar trend for Fe_d-p_ (crystalline and nano-crystalline minerals) (Fig. S4f). Exchangeable Ca and Mg exhibited consistent levels irrespective of depth (Fig. S4g, h), although Ca_ex_ content was markedly higher at the downstream location. Calcium carbonate, a primary component of shells and corals, is generally resistant to dissolution in a neutral pH (∼6.5) ammonium acetate solution. Thus, this exceptionally high downstream Ca_ex_ content was likely due to Ca ions retained in the soil as a result of long-term weathering of shell and coral deposits.

Among the extractable metals analyzed, clear positive correlations were found only between the OC content of HF (OC_HF_) and Al_p_ (R^2^=0.54, p=0.0018) and Fe_p_ (R^2^=0.87, p=5.1×10^-7^) (Fig. 5). This indicates that the dominant mode of organo-mineral or organo-metal associations in this mangrove soil were co-precipitation of OC and metals rather than sorption onto mineral surfaces (Fig. 5). Further supporting this, the OC-to-Fe_d_ mass ratios in HF were significantly higher than the maximum sorptive capacity of Fe oxide phases (Kaiser & Guggenberger, 2007) (Fig. 6). The strong correlation between OC_HF_ and Fe_p_ agreed with our hypothesis, but the relatively good correlation between OC_HF_ and Al_p_ suggested a similar role of Al and Fe in OC coprecipitation in this mangrove soil (Fig. 5b, e).

**Fig. 5.**
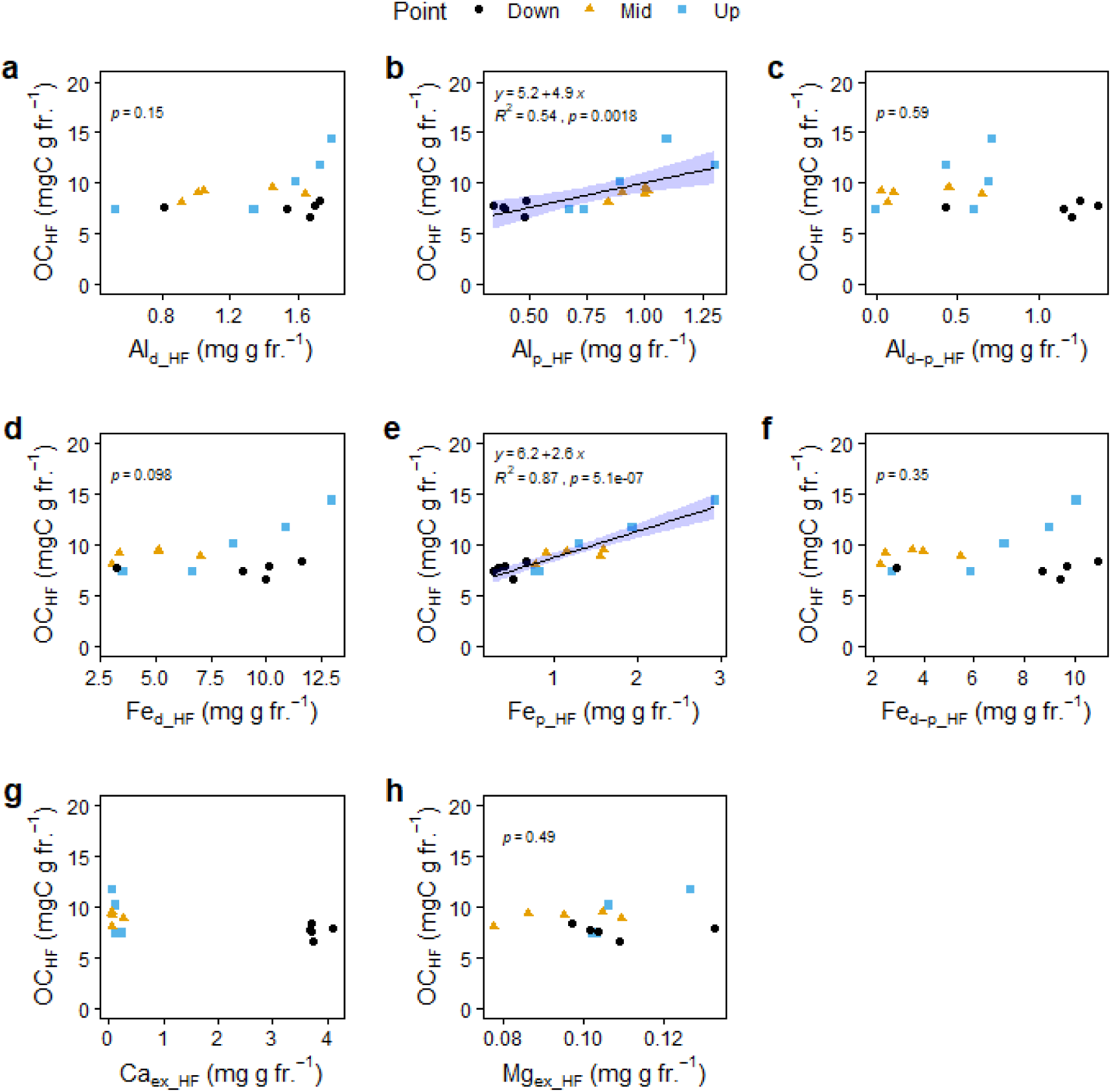
The correlations between extractable metals and organic carbon (OC) content in high-density fraction (HF). The shaded areas in (b) and (e) indicate the 95% confidence intervals. The subscripts represent dithionite-citrate extraction (*d*), sodium pyrophosphate extraction (*p*), and their differences (*d-p*), respectively. Ca_ex_ and Mg_ex_ are exchangeable calcium and magnesium, respectively. Correlation was not considered in (g) due to outliers.

**Fig. 6.**
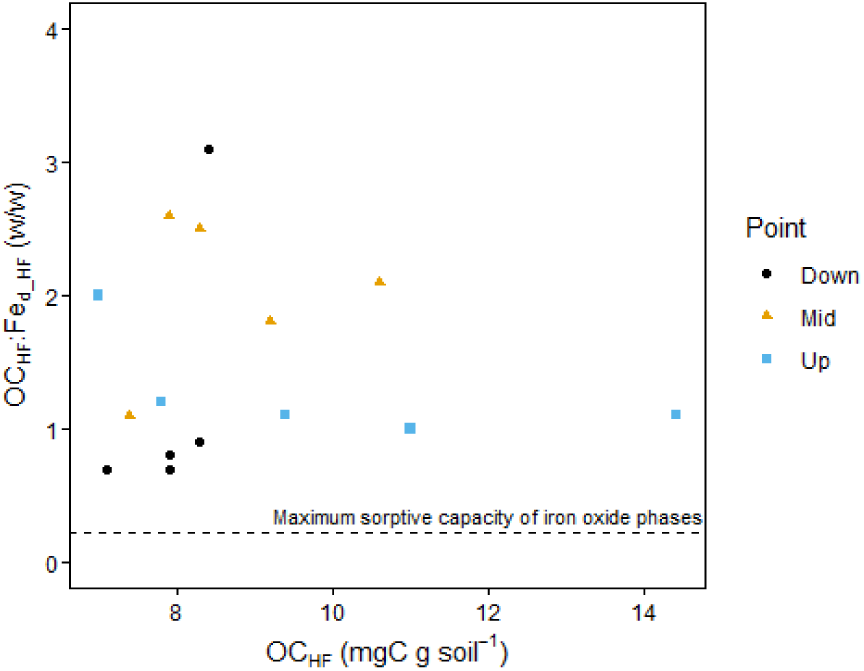
The mass ratio of OC to Fe_d_ in high-density fraction (HF) plotted against OC content in HF. The horizontal line represents the maximum sorptive capacity of FeOx phases: 0.22 g OC g Fe^-1^ = 0.8 mm^3^ OM (mm^3^ FeOOH)^-1^ from sorption experiments (Kaiser and Guggenberger, 2007).

### 3.5 Source plot

Scatter plots of C/N ratios, δ^13^C values, and Δ^14^C values of density fractions were analyzed and compared with those of endmembers to gain insights into possible OC sources in different fractions (Fig. 7). The most pronounced differences among fractions were observed in their C/N ratios (Fig. 7a). Compared to differences among fractions, differences by locations were much smaller and thus will not be discussed further. Based on C/N ratios and δ^13^C values, HF showed the closest similarity to terrestrial and riverbank soils and riverine SS rather than to LFs, the latter were considered primarily derived from mangroves. Contribution from marine sources was negligible in all samples (Fig. 7a). Δ^14^C values mostly separated HF from LFs (Fig. 7b, c).

**Fig. 7.**
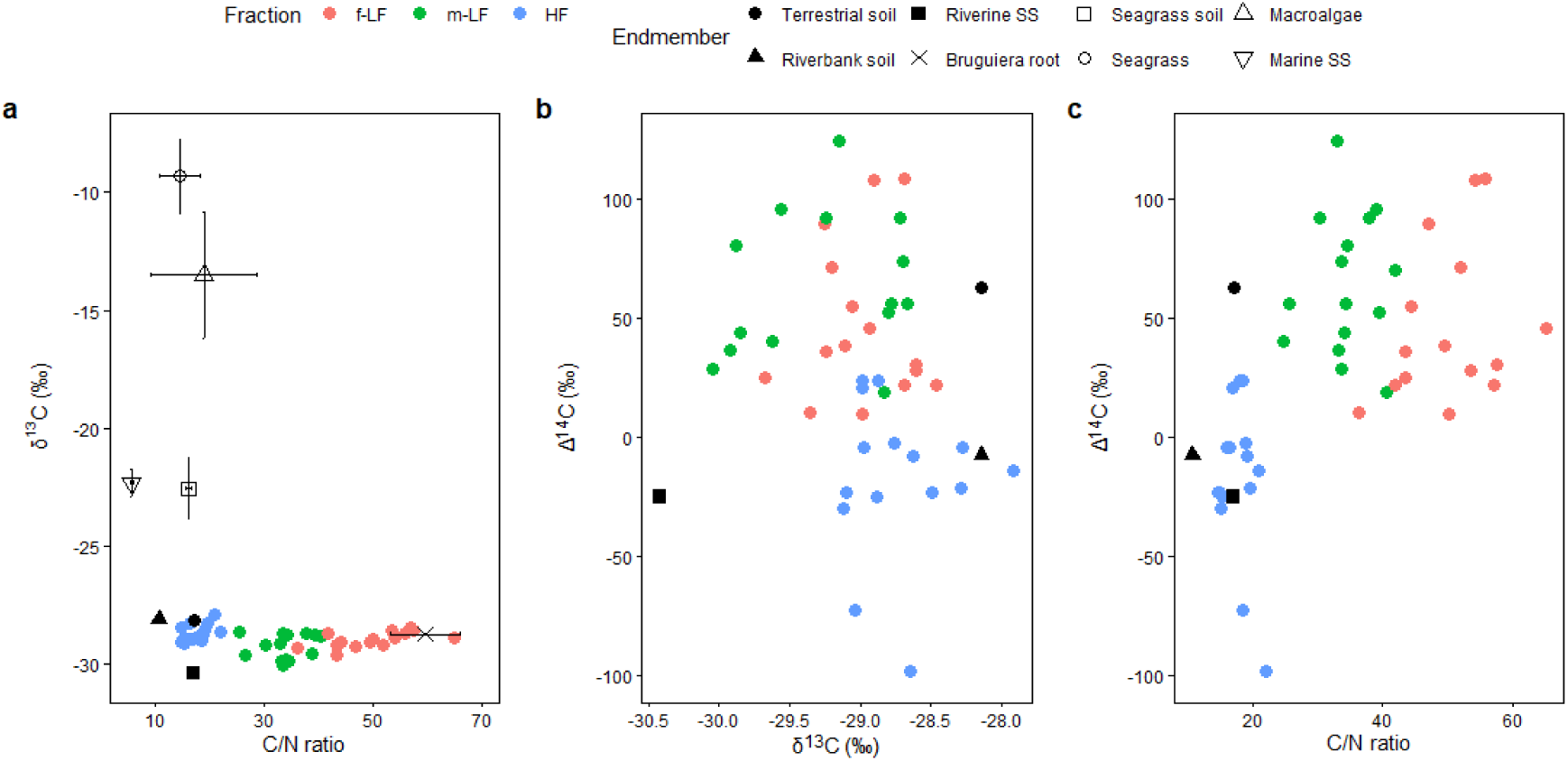
Scatter plots of C/N mass ratio, δ^13^C, and Δ^14^C values of density fractions and endmembers. SS = suspended solid. (a) Error bars represent the standard deviations, while endmembers with filled marks lack field replicates.

## 4. Discussion

Overall, the general characteristics of soil density fractions in the Gaburumata mangrove, such as the dominance of HF and abundant LFs throughout the core, align with findings from the nearby Fukido mangrove (Hamada et al., 2024), suggesting that these OC properties may be universal in sandy mangrove soils in the Subtropics. Discussions regarding fraction abundance, characterization, and relations to roots as well as extractable metals are discussed in Supplementary Information (S3).

### 4.1 The origin of organic matter in soil density fractions

The initial interpretation based on the commonly used C/N–δ^13^C plot suggested a clear shift in OC source with increasing mineral association (i.e., f-LF, over m-LF to HF) from mangrove roots to terrestrial OC, such as terrestrial and riverbank soils and riverine SS (Fig. 7a). However, source partitioning of OC is feasible only when the indicators of endmembers exhibit distinct values that remain relatively stable during degradation in the environment. Actually, the C/N ratio of plant residues can considerably decrease due to selective C leaching during initial decomposition and N retention through microbial reworking. Terrestrial endmembers, especially riverine SS, represent OC that has already undergone leaching and microbial decomposition prior to its deposition in mangrove soil. If C/N ratio of mangrove material remained constant while degradation as was assumed in past research (e.g., Fu et al., 2024), terrestrial OC would be a major contributor to m-LF and particularly HF (Fig. 7a).

However, morphological features and apparent radiocarbon age (Fig. 3c, d) strongly suggested that m-LF was a decomposed fragment of f-LF, which itself was predominantly derived from mangroves (Hamada et al., 2024). HF presumably forms through multiple processes, including the binding or inclusion of microbial-processed material from LFs with soil minerals and the sorption of leachates directly from LFs onto minerals (Lavallee et al., 2020).

The observed C/N ratios in each density fraction followed the sequence HF < m-LF < f-LF (Fig. 3a), with the lower C/N ratio of HF indicating more advanced microbial reworking of plant-derived material, consistent with terrestrial soil observations (Liao et al., 2006; Tan et al., 2007). Decreasing C/N ratios and increasing δ^13^C and δ^15^N values in heavier fractions were also reported in a study that used a series of heavy solutions to seagrass meadow soils (Miyajima et al., 2017). These results align with the well-established paradigm in soil science that OC_HF_ (or MAOM) originates predominantly from microbial-reworked, plant-derived material (Angst et al., 2021; Cotrufo et al., 2013; Lavallee et al., 2020). Therefore, we could not exclude the possibility that OC_HF_ was mostly mangrove derived. While diagenetic fluctuations in source indicators can be incorporated into Bayesian source mixing models (Parnell et al., 2013; Stock et al., 2018), changes in C/N ratios for mangrove material may be much more pronounced than previously considered. Based on differences in the C/N ratio (Fig. 7c) and age (Fig. 4) of corresponding f-LF and m-LF samples, the median annual change in C/N ratio was estimated to be 3 yr^-1^, meaning that C/N ratio can decrease more than 10 units in just a few years. Density fractionation followed by elemental and isotopic (δ^13^C, Δ^14^C) analyses will help evaluate this process under filed conditions. Finally, as δ^13^C remained almost constant among different density fractions and thus degradation, the clear distinction in δ^13^C values between mangrove and marine endmembers suggests a minimal contribution of marine sources to mangrove OC (Fig. 7a).

### 4.2. Organo-metal coprecipitates contributing to OC stabilization in mangrove mineral soil

The strong positive correlation between the Al_p_ and Fe_p_ contents and the OC content of HF (Fig. 5b, e), together with the high OC_HF_:Fe_d_ ratios (Fig. 6), suggested that organo-Al/Fe coprecipitation, rather than the sorption of OC onto metal (oxyhydr)oxides, play a crucial role in the stabilization of OC in mangrove soils. This finding is consistent with previous studies on coastal sediments and mangrove soils (Dicen et al., 2018; Lalonde et al., 2012; Ruiz et al., 2024; Shields et al., 2016; Zhao et al., 2018). These organo-metal associations lead to physical encapsulation of OC, protecting it from microbial degradation (Boudot et al., 1989). Past studies employing δ^13^C analysis of Fe-bound OC have shown that Fe preferentially preserves terrestrial material over marine material (Dicen et al., 2018; Shields et al., 2016; Zhao et al., 2018), in line with the minimal contribution of marine sources to HF found in this study (Fig. 7a). Although Fe is susceptible to reductive dissolution (Thompson et al., 2006), it is likely that redox fluctuations in the mangrove soil promoted enhanced co-precipitation with OC upon oxidation (Keil & Mayer, 2014). The slope of the OC_HF_-metal_p_ relationship indicated that approximately 12 carbons were associated with 1 iron atom or 11 carbons per 1 aluminum, respectively (Fig. 5b, e). Iron and aluminum ions probably formed co-precipitates with OC, thus the OC_HF_-(Fe+Al)_p_ relationship was examined (Fig. S5). They showed a strong linear correlation with the slope of 6.3 (Fig. S5b). Assuming that the clear linear response of OC_HF_ was solely due to increases in Fe_p_ and Al_p_ (Fig. S5b), this OC_HF_:metal_p_ ratio (6.3) was at the lowest range observed in volcanic ash soils (19.7 ± 15.3 in A horizons and 12.1 ± 4.2 in buried A horizons, analyzed for bulk soil) (Shimada et al., 2022), for which OC-metal_p_ relationships have been particularly studied. The positive y-intercept in the OC_HF_-metal_p_ relationship suggested the OC associated with other inorganic matrices, such as more crystalline Fe oxides and aluminosilicate minerals (Fig. S5b). The positive deviation of each point from the intercept in Fig. S5b approximated OC associated with metal_p_, which accounted for on average 44 ± 10% of OC_HF_, underpinning the significance of organo-metal complexes.

The sandy soil texture of this mangrove forest may explain the lack of effect of Ca_ex_ and Mg_ex_ on OC_HF_ (Fig. 5g, h). Mangrove forests on Ishigaki Island typically have a sandy texture (sandy loam to sandy clay loam), with sand content ranging from 60%–70% (Kida et al., 2017; Kida, Kondo, et al., 2019). Although no formal soil texture measurements were conducted in this study, hand analysis and visual observation suggested that the sand content of the Gaburumata mangrove was higher than that of the sandy loam soil of the Fukido mangrove. The low clay content reduced the importance of Ca_ex_ and Mg_ex_ on OC_HF_, because these divalent cations mostly capture OC through cation bridging, linking negatively charged clay surfaces and organic matter (Rowley et al., 2017). Since sand particles have a low surface charge, they do not facilitate cation bridging. The low clay content could also explain the minor role of crystalline and nano-crystalline Al and Fe minerals (Fig. 5a, c, d, f), as these secondary minerals exist predominantly in the clay fraction.

### 4.3. Inclusion of mangrove-derived fresh carbon into HF

The mismatch between root distribution (Fig. 1d) and the age of young OC fractions (Fig. 4ab) could have two possible explanations: (1) Our fine root density data based on one-time sampling may not represent year-round belowground OC inputs. In fact, mangrove fine root growth exhibits marked seasonality (Muhammad-Nor et al., 2019; Poungparn et al., 2015), and root exudate production likely shows similarly large seasonal variation, though studies on this are currently lacking. A more frequent, year-round evaluation of belowground fine root dynamics is clearly necessary. (2) Secondly, and maybe more importantly, fine root decomposition may occur at different rates depending on depth. The faster turnover of f-LF and m-LF at surface layers compared to mid depths (Fig. 4ab) may be possible by presumably more oxidized conditions that favor oxic litter decomposition (Kida & Fujitake, 2020) . The fast turnover of f-LF and m-LF at the deepest depth (Fig. 4ab) is more difficult to explain but may result from preferential water flow at depth by animal burrows creating favorable decomposition conditions (Y. Hua et al., 2024; Tomotsune et al., 2019; Yin et al., 2023) or the abundant production of fine roots at depth (Arnaud et al., 2021; Fujimoto, 2021; Ohtsuka et al., 2024; Ono et al., 2022). The similar depth distribution of LF ages across sampling locations suggests that mechanisms influencing the apparent LF turnover were universal within this mangrove forest (Fig. 4ab).

The Δ^14^C–OC relationship suggested an efficient incorporation of mangrove-derived modern C into HF, in addition to the pre-existing old C (Fig. 3d), against general understanding of HF as having a slow turnover time (Heckman et al., 2021). Nevertheless, several studies have reported a rapid response of HF to plant inputs in terrestrial soils (Koarashi et al., 2012; Schrumpf et al., 2013; Schrumpf & Kaiser, 2015; Swanston et al., 2005). For instance, Swanston et al. (2005) found rapid (a few years) incorporation of ^14^C into HF in stand-level ^14^C-labeling in oak forests. Similarly, Koarashi et al. (2012) estimated that a significant proportion (28–73%) of OC_HF_ turned over on the time scales of decades in the subsurface layers of grassland and forest soils, despite their apparent old ages. Although the age depth distribution was opposite between HF and LFs (Fig. 4), this could be explained by possible differences in LF decomposition by depth, as explained previously. Enhanced mineralization of LF at surface or deep sections would result in less incorporation of mangrove-derived modern carbon to HF, while longer turnover of LF at mid depths could contribute to incorporation of modern carbon to HF, making the apparent age of HF younger (Fig. 4c).

Our observation has two key implications: first, OC in the mangrove forest soil consisted of sources with distinct ^14^C ages, and samples with lower OC_HF_ content and more depleted ^14^C had a lower contribution from modern mangrove-derived C. This aligns with findings from salt marsh sediments, where OC consisted of old allochthonous (terrestrial) and young autochthonous (marsh-derived) C (Broek et al., 2018; Komada et al., 2022). Kamada et al. (2022) estimated that less than half of the OC in the meso-density fraction (1.6 < density < 2.5 g cm^-3^) in salt marsh sediments was derived from autochthonous production. In the present study, the limited sample size and the presence of outliers in the Δ^14^C–OC relationship (Fig. 3d) prevented estimation of mangrove-derived C using a mixing and net-reaction model (Komada et al., 2022). Nevertheless, our results suggest that radiocarbon analysis offers a more reliable means of source attribution for HF in mangrove soils, a challenge when source indicators overlap between multiple endmembers (e.g., δ^13^C values of C3 terrestrial plants and mangroves) or change by decomposition (e.g., C/N ratio) (Fig. 7a).

Second, mangrove expansion in subtropical and tropical regions, driven by global warming and plantation efforts, may enhance coastal stable OC stocks due to rapid inclusion of mangrove-derived OC into soil mineral matrix (i.e., HF). Warming has facilitated mangrove encroachment into tidal flats and salt marshes globally (Kelleway et al., 2017), with studies showing increased OC stocks following mangrove expansion or plantation (Doughty et al., 2016; Ray et al., 2023). However, the form of OC newly accumulated after mangrove expansion, whether free or mineral-associated, remains poorly understood. Our results suggest that mangrove expansion may quickly increase stable, mineral-associated OC. Furthermore, given that more than half of existing OC stock in mangroves with mineral soils are mineral associated (G. Chen et al., 2023; Fu et al., 2024; Hamada et al., 2024), the temperature sensitivity of this fraction (i.e., OC_HF_) is critical. A meta-analysis of warming studies in terrestrial soils suggests that increasing temperature will not reduce OC_HF_ (Rocci et al., 2021), as its decomposition is restricted by the inaccessibility of microbes and enzymes, rather than their temperature-dependent activity (Gentsch et al., 2018). Lesser temperature sensitivity of OC_HF_ in the global compilation of LF and HF content in terrestrial soils also supports the result of warming studies (Georgiou et al., 2024). Collectively, our analysis suggests that mangrove-derived modern carbon is efficiently incorporated into the mineral-associated fraction, offering an optimistic perspective that mangrove expansion enhances stable soil OC pools in addition to increasing plant biomass and litter (Ohtsuka et al., 2024). This accumulation of stable OC pool can partially offset the accelerated decomposition of non-protected plant litter under warming conditions (Arnaud et al., 2020).

### 4.4. Limitations and future opportunities

#### Spatiotemporal sampling

The importance of organo-metal coprecipitation in OC_HF_, as highlighted in this study (Fig. 5b, e), also raised the necessity of finer-scale spatiotemporal sampling. Organo-Fe precipitation, which occurs rapidly and is potentially reversible (Thompson et al., 2006), presumably exhibits highly dynamic behavior in redox-sensitive environments such as mangrove soils. These soils are subject to daily, lunar, and seasonal tidal fluctuations that alter redox conditions, further complicated by localized oxygen intrusion from animal burrows and mangrove roots (Cheng et al., 2012; Kristensen & Alongi, 2006).

Differences in the redox sensitivity of metals can also influence the association of specific metals with OC. For instance, Nitzsche et al. (2022) found that under more acidic and anoxic conditions, Al preferentially binds with less-microbially processed OC compared to Fe (Nitzsche et al., 2022). While we partially addressed sample heterogeneity by using composite samples (see *Methods*), on-site temporal sampling and finer-scale gridded sampling could have provided additional insights. Sampling of soils around mangrove roots (i.e., rhizosphere) is potentially interesting to target Fe that accumulates around mangrove roots due to oxygen release from roots.

#### Soil texture

The negligible role of Ca_ex_ and Mg_ex_ in influencing OC_HF_ may be site-specific to the sandy soil of the studied mangrove forest (Fig. 5g, h). These cations could play a more significant role in clay-rich soils that provide the matrix for cation bridging. Studies encompassing different soil textures could provide a broader picture of relationships between soil texture and influential factors on OC accumulation.

#### Source partitioning

Although we showed the potential of radiocarbon analysis in improving source partitioning of OC in mangrove soils, further improvements may be achieved through biomarker analyses specific to OC sources (Resmi et al., 2016). Particularly challenging is the separation of mangrove-derived OC from that of other C3 plants within the catchment (Fig. 7). Physical separation of OC fractions based on their decomposition states and mineral associations seems prerequisite for meaningful source attribution for this heterogeneous mixture.

## 5. Conclusions

We applied density fractionation to investigate the mechanisms underlying OC preservation in mangrove soils. Reactive metal phase analysis highlighted the important role of organo-Al/Fe co-precipitation in stabilizing OC within mangrove soils. The mineral-associated OC (high-density fraction) constituted more than half of the total OC stock in the studied mangrove forest, with 44 ± 10% of this mineral-associated OC likely stabilized through organo-metal coprecipitation. Radiocarbon evidence further indicated efficient incorporation of modern, mangrove-derived carbon to mineral-associated OC. As mineral-associated OC is generally more refractory than particulate OC and potentially insensitive to temperature-driven decomposition, mangrove expansion is likely to enhance stable soil OC pools alongside increases in plant biomass and litter. This study offers an optimistic perspective on OC storage in mangrove soils under global warming and underscores the need for future research to explore the forms and stability of OC newly accumulated following mangrove expansion, which can be investigated through, for instance, chronosequence studies.

## Author contributions

KH collected samples and conducted experiments with the help from MK. KH conducted stable isotope analysis with the help of TO. NY conducted root density analysis, while TM collected and analyzed seagrass, macroalgae, and marine suspended solids. YM and YY conducted radiocarbon analysis. KH wrote an initial draft with a significant contribution from MK, and all authors have reviewed and approved the final article.

## CRediT Classification

Conceptualization (MK), Data Curation (MK), Formal Analysis (KH, MK, NY, TM), Funding Acquisition (MK, TO, TM, YY), Investigation (KH, MK, NY, TM, YY, YM), Project Administration (MK, TO, NF), Resources (MK, NF, TM, YY), Software (KH, MK), Supervision (MK), Visualization (KH, MK), Writing – Original Draft Preparation (KH, MK), Writing – Review & Editing (all authors).

## Data availability

Original data can be found in open-access repository PANGAEA at doi: .

## Acknowledgements

This study was supported by JSPS KAKENHI Grant Numbers 24H01513 (MK), 21KK0186 (TO), 20H00193 (YY), and 22H03717 (TM) and CREST JPMJCR23J6 (YY). MK was also partially supported by Tenure-Track Fellowship of Kobe University Institute for Advanced Research. We thank Yuta Watanabe, Ayaka Nishikawa, and Masahiro Shimotani from Kobe University for their help in extractable metal analysis. We thank collaborators and students from Gifu University, University of the Ryukyus, Waseda University, and The University of Shiga Prefecture, for their assistance in the field survey which resulted in Fig. 1c.

