## Supporting Information for "Organo-metal coprecipitation contributes to stable organic carbon fraction in mangrove soil" (from another version of this preprint: version 1, 2025-02-19)

**Contents:**

Supplementary method (S1, S2)

Supplementary discussion (S2)

Five Supplementary figures

Supplementary references

### S1. Optimization of the experimental condition for density fractionation

Density fractionation was performed using an aqueous solution of sodium polytungstate (SPT-0, TC-Tungsten Compounds, Germany; SPT) with a density of 1.6 g $cm^{-3}$, following established methodologies (Golchin et al., 1994; Hamada et al., 2024; Wagai et al., 2008). This method, widely used in terrestrial soils to separate soil organic matter fractions of varying microbial degradability, has seen limited application in mangrove soils (Hamada et al., 2024). Given the unique properties of wet soils such as mangroves, optimizing density fractionation conditions is crucial. In this study, we investigated the effects of soil sample conditions (field-moist or freeze-dried) on density fractionation and appropriate ultrasonic output during m-LF fractionation operations, conditions we could not check in a previous study (Hamada et al., 2024). Another set of soil samples was collected at five different depths (10-20 cm, 30-40 cm, 50-60 cm, 70-80 cm, and 90-100 cm) from the Gaburumata River for this purpose and divided into field-moist and freeze-dried samples. Field-moist samples were sieved with a 4-mm mesh sieve, while freeze-dried samples were sieved with a 2-mm mesh sieve. Subsequently, the density fractionation method, following a previously established protocol for mangrove soils (Hamada et al., 2024), was applied to both sets of samples, and the weight of each density fraction was determined. To account for water content, 10 subsamples of field-moist samples were dried at 105°C for 24 hours to determine moisture content, which was then used to calculate the recovered weight of density fractions (Fig. S1). Although no significant differences were observed in the relative abundance of each fraction between the two treatments and the distributional trends along depth were very similar (Fig. S1abc), the weight recovery of field-moist soils was 5 to 10 percentage points lower than that of freeze-dried samples (Fig. S1d). This lower recovery was attributed to potential errors in estimating water content using different subsamples rather than material loss during density fractionation. Therefore, field-moist samples pose challenges to density fractionation, including errors in moisture content estimation and the need for more careful subsampling to obtain homogeneous, reproducible samples. In contrast, freeze-dried soils exhibit excellent homogeneity and preservability and were used for subsequent analyses.

We then assessed the influence of ultrasonic power on the recovery of mineral-associated light fraction (m-LF) from sandy mangrove soils. Previous studies have highlighted the significance of ultrasonic conditions, particularly the total energy and power, in ensuring effective dispersion of OC fractions (Hamada et al., 2024; Poeplau & Don, 2014). Hamada et al. (2024) have shown that sonication at a total energy of 120 J $mL^{-1}$ effectively recovered the m-LF fraction. In this study, we kept the total energy constant at 120 J $mL^{-1}$ while varying ultrasonic power levels (20 W, 40 W, 60 W, 80 W) to investigate their impact on m-LF recovery. The m-LF recovery across different power settings was small and practically negligible compared to field data reported from the Fukido mangrove (Hamada et al., 2024) (Fig. S2), indicating that the m-LF likely originated from easily disintegrating aggregates. However, the time required for collecting and rinsing m-LF varied significantly with power output. Rinsing at lower power (20 W) took approximately 2 hours per sample, whereas at higher power (80 W), it extended to about 6 hours due to membrane filter clogging caused by finer m-LF fragmentation. Considering these factors, processing at 20 W proved more efficient, leading us to conduct subsequent density fractionation under these conditions (120 J $mL^{-1}$, 20 W, freeze-dried soils).

### S2. Details of sample analysis

#### S2.1 Inorganic carbon removal by HCl fumigation

The IC removal process was carried out on seagrass and mangrove soil samples (bulk soil and HF). Briefly, we encapsulated finely ground soil samples in Ag capsules (9 × 5 mm, LUDISWISS, Switzerland) and placed them in a glass 96-well microplate. We added ultrapure water by a solid-liquid mass ratio of approximately 1:1 to accelerate HCl vapor contacting samples. We placed a glass petri dish containing about 15 mL of 12 M HCl at the base of a clean, acid-washed glass desiccator, with the microplate containing the capsules placed on top. To enhance vaporization of concentrated HCl, we reduced the pressure inside the desiccator by a vacuum pump and exposed the samples to HCl for 6 hours. After exposure, we dried the microplate and samples at 105°C for 12 hours. Due to the fragility of the Ag capsules post-exposure, we sealed them in tin capsules (8 × 5 mm, Elemental Microanalysis, England) for elemental analysis.

#### S2.2 Extractable metals analysis

Briefly, each extraction of Al and Fe (sodium dithionite-citrate extraction and sodium pyrophosphate extraction) involved placing 0.2 g of finely ground HF sample in a 50 mL polyethylene bottle with a solid-to-liquid ratio of 1:100, followed by reciprocal shaking at room temperature for 16 hours. For dithionite-citrate extraction, fresh dithionite powder (0.2 g, technical grade 85%, Sigma-Aldrich) was added to each sample. After extraction, we transferred approximately 10 mL of the resulting solution to a 15 mL centrifuge tube and treated with a drop of 4 g $L^{-1}$ nonionic polyacrylamide to aid in flocculation. We isolated the supernatant by centrifugation at 10000 G for 10 minutes (dithionite-citrate extraction) or 26200 G for 30 minutes (sodium pyrophosphate extraction). Next, we added an yttrium standard solution (250 ppm) to the supernatant, which was then diluted 15-fold (dithionite-citrate extraction) or 3-fold (sodium pyrophosphate extraction) using ion-exchanged water.

We extracted exchangeable Ca ($Ca_{ex}$) and Mg ($Mg_{ex}$) using a modified batch method involving 0.05 M ammonium acetate and 0.0114 M strontium chloride ($SrCl_2$), a technique widely applied in Japan (Kamewada & Shibata, 1997; Matsue & Wada, 1985). This quick method improves Ca extraction efficiency due to strong affinity of Sr for soil matrix, yielding comparable amounts of Ca, Mg, and potassium to those obtained via the original Schollenberger column method. Briefly, we weighed 0.2 g of finely ground HF sample in a 50 mL polyethylene bottle with a solid-to-liquid ratio of 1:200 and shook it reciprocally at room temperature for 1 hour. After extraction, we separated the supernatant by centrifugation at 10000 G for 10 minutes, spiked it with 250 ppm yttrium, and analyzed it by ICP-AES employing the internal standard method with calibration curves matching the sample matrix

**S3. General characteristics of mangrove soil density fractions and extractable metals**

The abundant LFs until a depth of 1 m aligned with massive fine root distributions at depth (Figs. 1d, 2a). Recent studies have reported that fine root production predominantly occurs at depths exceeding 30 cm (Arnaud et al., 2021), with some also extending to 50–100 cm deep (Fujimoto, 2021; Ohtsuka et al., 2024; Ono et al., 2022). Our results were consistent with these findings, underscoring the importance of fine roots as belowground OC sources. The dominance of dead fine roots suggested rapid turnover of fine roots (Fig. 1d). Thus, understanding the decomposability of dead fine roots and their potential incorporation into the soil matrix as stable soil OC is crucial for OC dynamics in mangrove forests, a topic we discuss in section 4.4. There was a pronounced decrease in N contents in LFs with depth (Fig. S3c), suggesting selective preservation of C-rich compounds such as phenolic materials (Kida et al., 2019), or differences in the activity of N-fixing bacteria by depth (Luo et al., 2021) .

HF was the most abundant among density fractions, while both f-LF and m-LF increased with depth across locations (Fig. 2a). In HF, a clear increasing trend with depth was observed only at the upstream location (Fig. 2a), reflecting the increase in C and N contents on fraction basis (Fig. S3c). Other studies have reported high variability in density fraction abundance in mangrove soils (Hamada et al., 2024) or declines in POM and MAOM abundance with depth in mangrove and salt marsh soil (Fu et al., 2024). On a N basis, HF was approximately five times more abundant than LFs (Fig. 2b), and the relative contribution of each fraction showed minimal variation with depth. In line with findings from the nearby Fukido mangrove (Hamada et al., 2024), m-LF contributed approximately 1.4 times more than the average terrestrial soil to bulk OC (Heckman et al., 2021). In mangrove soils, the prolonged residence time of plant residues, predominantly in the form of fine roots or f-LF, under suboxic conditions likely enhanced associations with soil minerals. The concurrent increases in C/N ratios and $\delta^{13}C$ values with depth across all fractions suggested a higher relative contribution of roots over leaves (Fig. 3a, b), as previously observed in the Fukido mangrove (Hamada et al., 2024). Overall, these general characteristics of soil density fractions align with findings from the nearby Fukido mangrove (Hamada et al., 2024), suggesting that these OC properties may be universal in sandy mangrove soils in the Subtropics.

The strong correlation between $Al_d$ and $Fe_d$ contents ($r = 0.89$, $p < 0.001$) (Fig. S4a, d) suggests that their distributions were likely influenced by common processes, such as deposition of SS containing both elements and co-precipitation of Al with Fe(III) hydroxides upon oxidation, despite their differing susceptibility to redox conditions (Nitzsche et al., 2022). The $Al_d$ and $Fe_d$ contents increased with depth (Fig. S4), consistent with mangrove soils in the Philippines (Dicen et al., 2018), while contrasted with coastal wetland sediments and mangrove soils in China where $Fe_d$ content decreased with depth (Li et al., 2024; Sun et al., 2023). Available data on reactive metals in similar environments is limited, preventing making general conclusions.

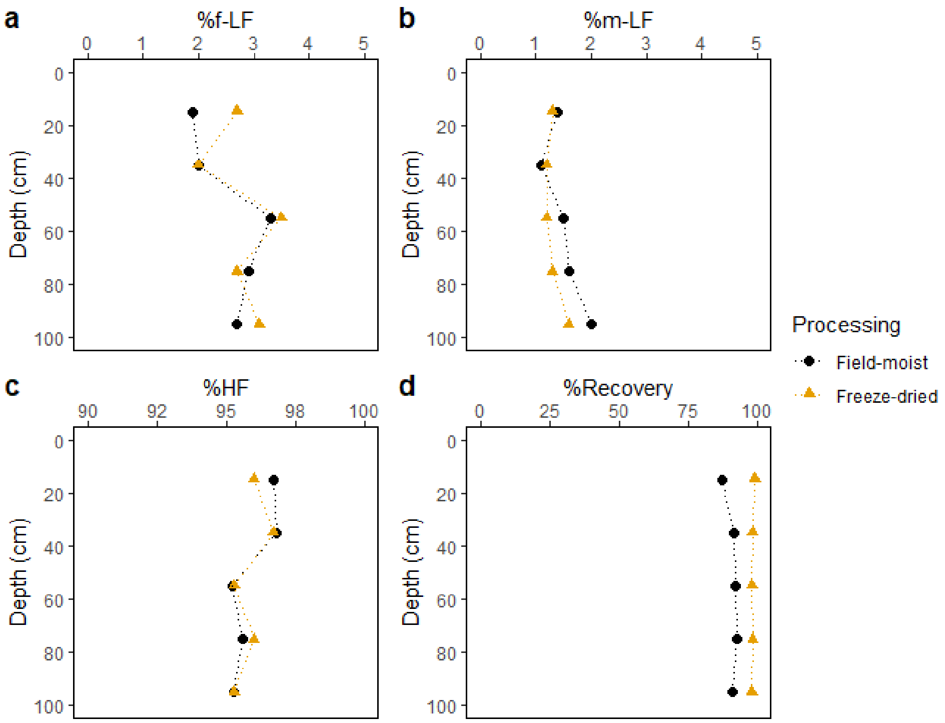


Fig. S1. The weight proportions of each fraction in field-moist and freeze-dried samples with depth.

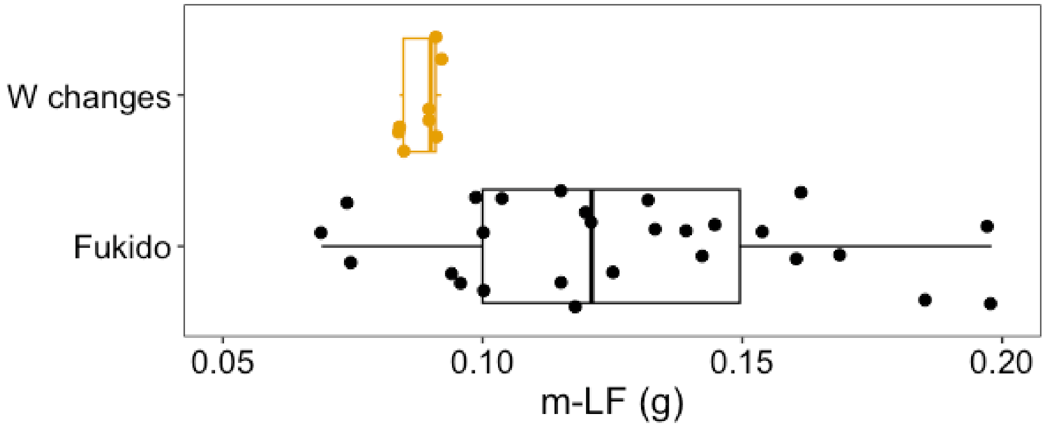


Fig. S2. The m-LF recovery weights measured under varying ultrasonic power outputs (20, 40, 60, 80W), at a constant energy level of 120 J mL$^{-1}$. The results for "Fukido" illustrates the m-LF recovery in nearby Fukido mangrove forest soils (Hamada et al., 2024). m-LF: mineral-associated low-density fraction.

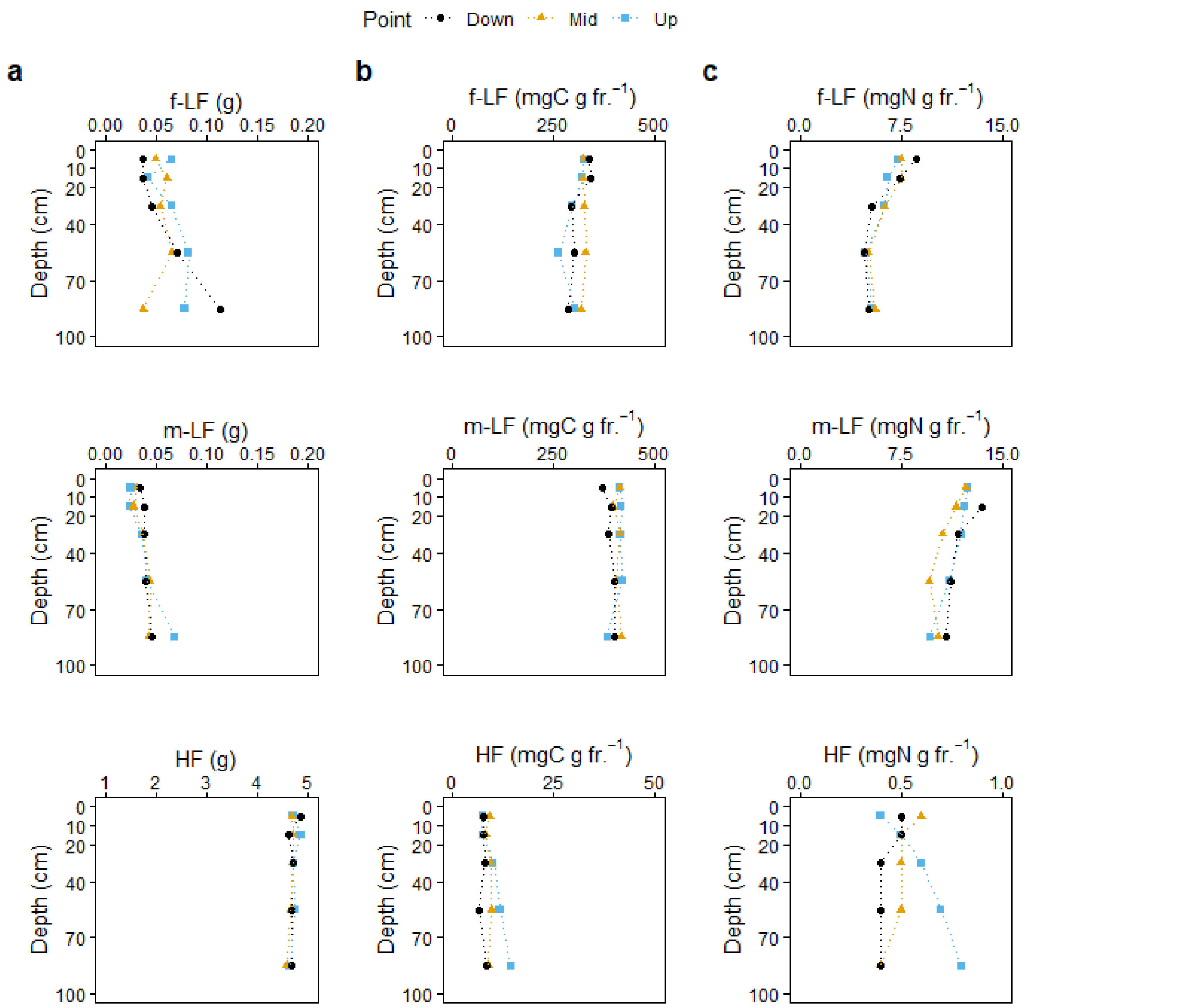


Fig. S3. Contribution of each density fraction to total soil mass (partitioned from 5 g) (a) and C and N concentrations of each fraction on fraction basis (b, c) with depth. f-LF: free low-density fraction, m-LF: mineral-associated low-density fraction, and HF: high-density fraction.

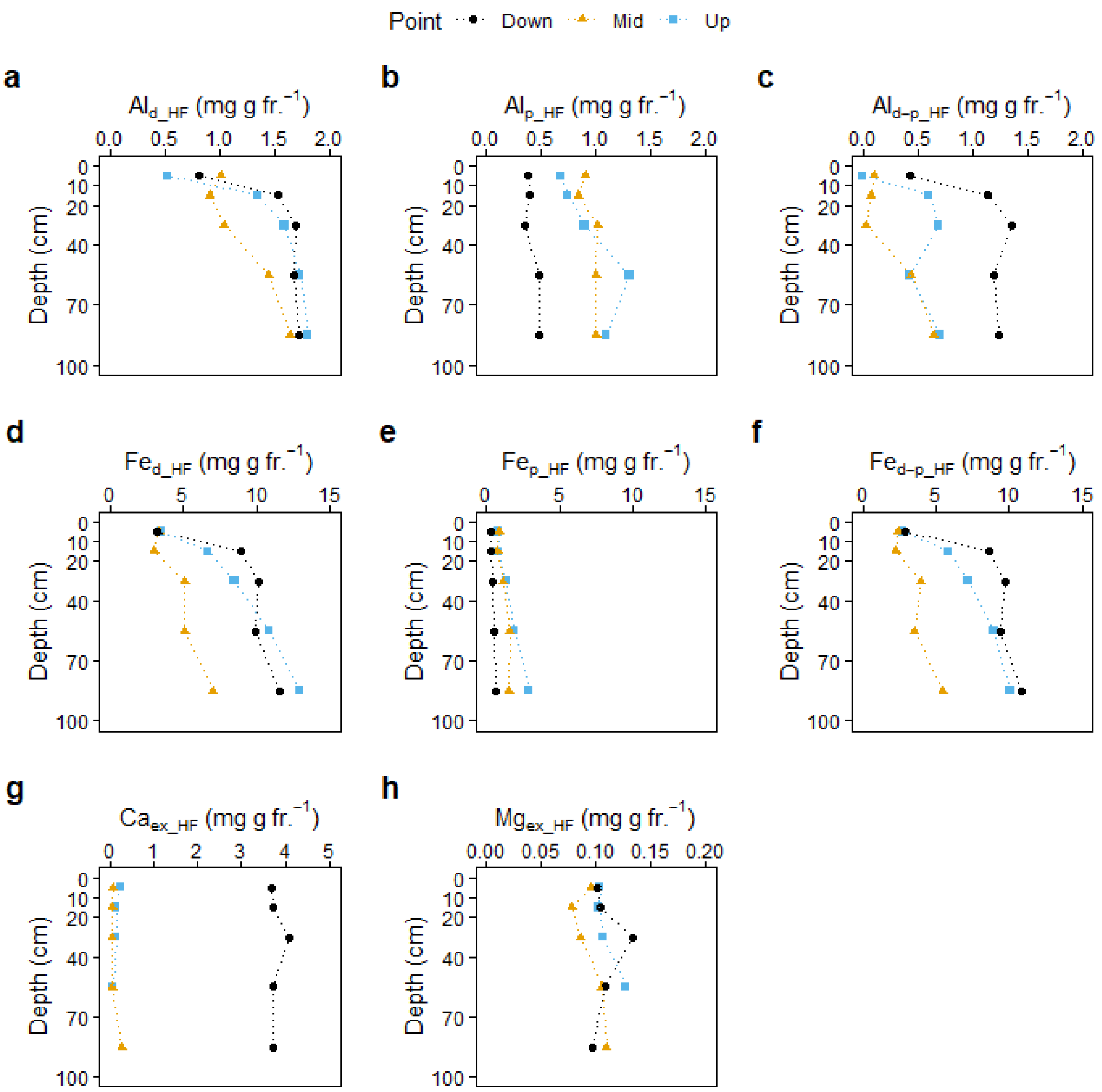


Fig. S4. Depth profiles of extractable metals in high-density fraction (HF). The subscripts indicate dithionite-citrate extraction (*d*), sodium pyrophosphate extraction (*p*), and their differences (*d-p*), respectively. $Ca_{ex}$ and $Mg_{ex}$ are exchangeable calcium and magnesium, respectively.

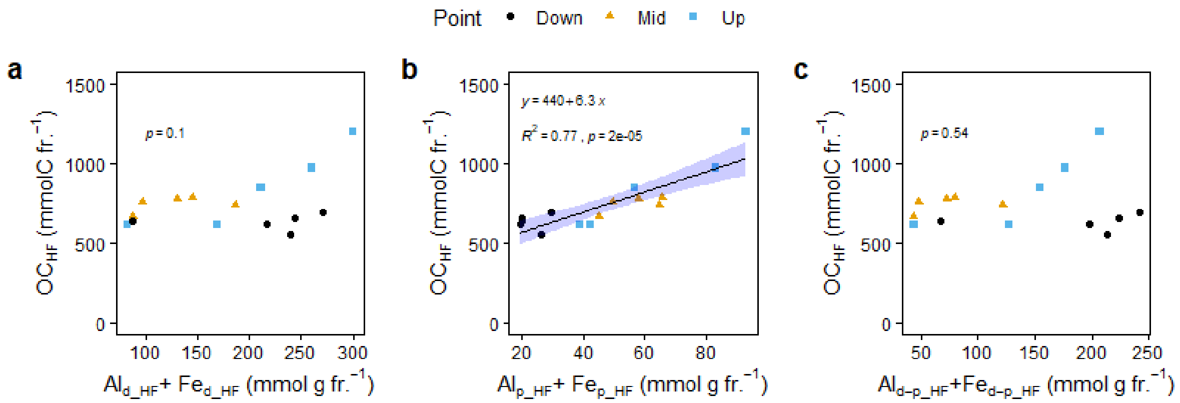


Fig. S5. The correlations between extractable metals (Al+Fe) and organic carbon (OC) content in high-density fraction (HF) on a molar basis. The shaded areas in (b) indicate the 95% confidence intervals. The subscripts represent dithionite-citrate extraction (*d*), sodium pyrophosphate extraction (*p*), and their differences (*d-p*), respectively.
